# High-Codon: A Deep Learning-Based Codon Optimization Tool for Enhanced Heterologous Protein Expression in Escherichia coli

**DOI:** 10.1101/2025.05.30.656984

**Authors:** Jiawei Li, Xinxiu Dong, Jie Liu

## Abstract

High-Codon is a deep learning-based codon optimization tool designed to enhance the expression levels of heterologous proteins in Escherichia coli. This approach employs a BERT pre-trained model to construct a sequence labeling framework for predicting optimal codons. Additionally, an expression-level weighted loss function is introduced to strengthen the model’s ability to learn from highly expressed proteins. Evaluation using 100 protein sequences from 34 species demonstrates that High-Codon outperforms traditional optimization methods.

## Introduction

The prokaryotic expression system is widely utilized in recombinant protein production, vaccine development, and antibody preparation due to its high efficiency, low cost, and ease of operation^1–3^. In these applications, the expression level of heterologous proteins is a critical factor determining their functional efficiency. Codon usage directly influences protein expression efficiency and yield, playing a pivotal role in optimizing protein production^4^.

A codon is a triplet of three consecutive nucleotides that determines the coding of amino acids during protein synthesis. In the transmission of genetic information, multiple codons can encode the same amino acid, and these codons are referred to as synonymous codons. However, the usage frequency of synonymous codons is not uniform during protein synthesis. Different organisms or genes often exhibit a preference for specific synonymous codons, a phenomenon known as codon usage bias^5^. The codons that are preferentially used are referred to as optimal codons^6^.

The fundamental principle of codon optimization is based on codon bias, which refers to the preferential usage of synonymous codons that varies among different species^7^. Codon usage bias is influenced by multiple factors, including mutational pressure and natural selection. The selection of different codons is closely related to tRNA availability, ribosome stalling, and mRNA secondary structure, among other factors. These complex regulatory mechanisms act in concert to determine protein expression levels^8^.

In heterologous protein expression systems, codon optimization is a crucial strategy to enhance the translation efficiency and protein expression levels of target genes in the host cell. According to the study by Claes Gustafsson et al., selecting synonymous codons that are more frequently used in the host genome can significantly improve heterologous expression, and the use of particular codons can increase the expression of a transgene by more than 1,000-fold^9^.Therefore, codon optimization plays a critical role in achieving high-level heterologous protein expression.

Due to the degeneracy of the genetic code, different codons can encode the same amino acid. Among the 20 standard amino acids, a total of 61 codons participate in encoding, leading to an enormous number of potential codon arrangements. Theoretically, for a polypeptide chain composed of 100 amino acids, the possible codon arrangements reach 10^78^ (i.e., 61^100^). Such an immense combinatorial space renders exhaustive search computationally infeasible, necessitating the development of efficient codon optimization algorithms.

Traditional codon optimization strategies are primarily based on statistical metrics, aiming to enhance expression efficiency by replacing rare codons with host-preferred ones. However, excessive reliance on high-frequency codons may lead to imbalances in the tRNA pool, ultimately resulting in tRNA depletion and premature translation termination^10^.

More importantly, such statistical approaches are inherently unable to capture deeper-level information, such as codon context dependencies, mRNA secondary structures, translation dynamics, and ribosome occupancy patterns. These factors play a crucial role in gene expression. For instance, codon arrangements can influence ribosome movement speed, thereby affecting co-translational protein folding, while local mRNA structures may impact translation initiation or elongation efficiency^11,12^. Therefore, relying solely on statistical methods for codon optimization often fails to achieve optimal expression outcomes.

Several recent studies have advanced codon optimization through deep learning-based approaches. For example, the method proposed by Hongguang Fu et al. integrates a BiLSTM-CRF model with codon boxes to improve the codon selection process^13^. The ICOR (Improving Codon Optimization with Recurrent Neural Networks) tool successfully captures complex codon usage patterns by training a BiLSTM model on large-scale genomic data, generating sequences that better reflect natural codon preferences^14^. Additionally, Haoran Gong et al. developed iDRO (integrated deep-learning-based mRNA optimization), which optimizes both the open reading frame (ORF) and untranslated regions (UTRs) of mRNA sequences. Their approach leverages a BiLSTM-CRF model for codon selection and an RNA-Bart transformer for UTR generation, resulting in optimized sequences that mimic human endogenous gene patterns and enhance protein expression^15^. Furthermore, Dennis R. Goulet et al. demonstrated the effectiveness of an RNN-based codon optimization model trained on Chinese hamster DNA sequences. Their model generated optimized DNA sequences that, when expressed in Chinese hamster ovary cells, produced protein levels comparable to or higher than those achieved with conventional optimization methods^16^. These advancements demonstrate that deep learning-based codon optimization strategies are feasible and hold great potential for future applications.

In previous studies, researchers often selected sequences with high Codon Adaptation Index^17^ (CAI) for training. The CAI is calculated using the following formula:

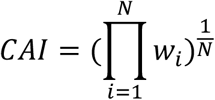

However, training models solely on sequences with high CAI scores may overlook lowly expressed genes across the entire genome, thereby missing negative samples and special cases, which could negatively impact the model’s generalizability.

Moreover, previous methods assigned equal weights to different sequences during training, disregarding the actual expression levels of different genes in the organism. This could potentially limit the model’s ability to enhance expression levels effectively.

High-Codon proposes a deep learning-based codon optimization method to enhance heterologous protein expression levels in Escherichia coli. Specifically, High-Codon utilizes a BERT-based pre-trained model for sequence labeling to predict the optimal codon for a given amino acid sequence. To further optimize the model, we introduce a weighted loss function that incorporates protein expression levels as an additional training signal, thereby enhancing the model’s ability to learn from highly expressed genes. The model performance is evaluated using codon adaptation index (CAI) and GC content.

## Results

**Figure 1.**
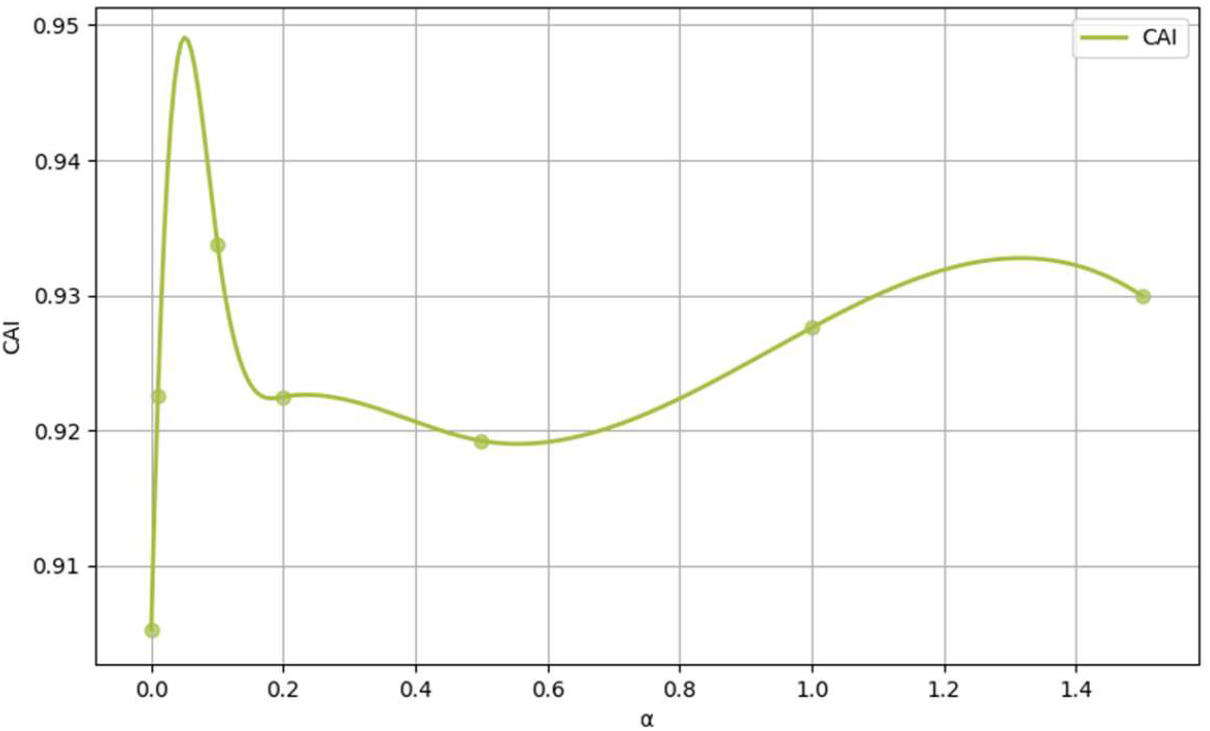

To evaluate the impact of expression-weighted training, we varied the scaling factor (α) and measured the Codon Adaptation Index (CAI) on the test set. As shown in Figure X, the CAI increased from 0.905 at (α= 0) (no weighting) to a peak of 0.934 at (α= 0.1), indicating that introducing expression-based weighting can enhance codon optimization. As α increased further, CAI slightly decreased to 0.919 at (α= 0.5), then gradually rose again, reaching 0.930 at (α= 1.5). This non-monotonic trend suggests that both overly small and overly large values of α lead to suboptimal performance, with moderate weighting yielding the best results.

Considering both CAI and other sequence-level properties such as GC content, we selected the model trained with α for downstream analysis.

### Performance Evaluation

To ensure that the model can be widely applied to the expression of different proteins in Escherichia coli, 100 protein sequences were selected randomly from 34 species and evaluated based on Codon Adaptation Index (CAI) and GC content.The performance of our model was compared with two widely used codon optimization tools, **GenSmart** and **ExpOptimizer**.

### CAI

**Figure 2.**
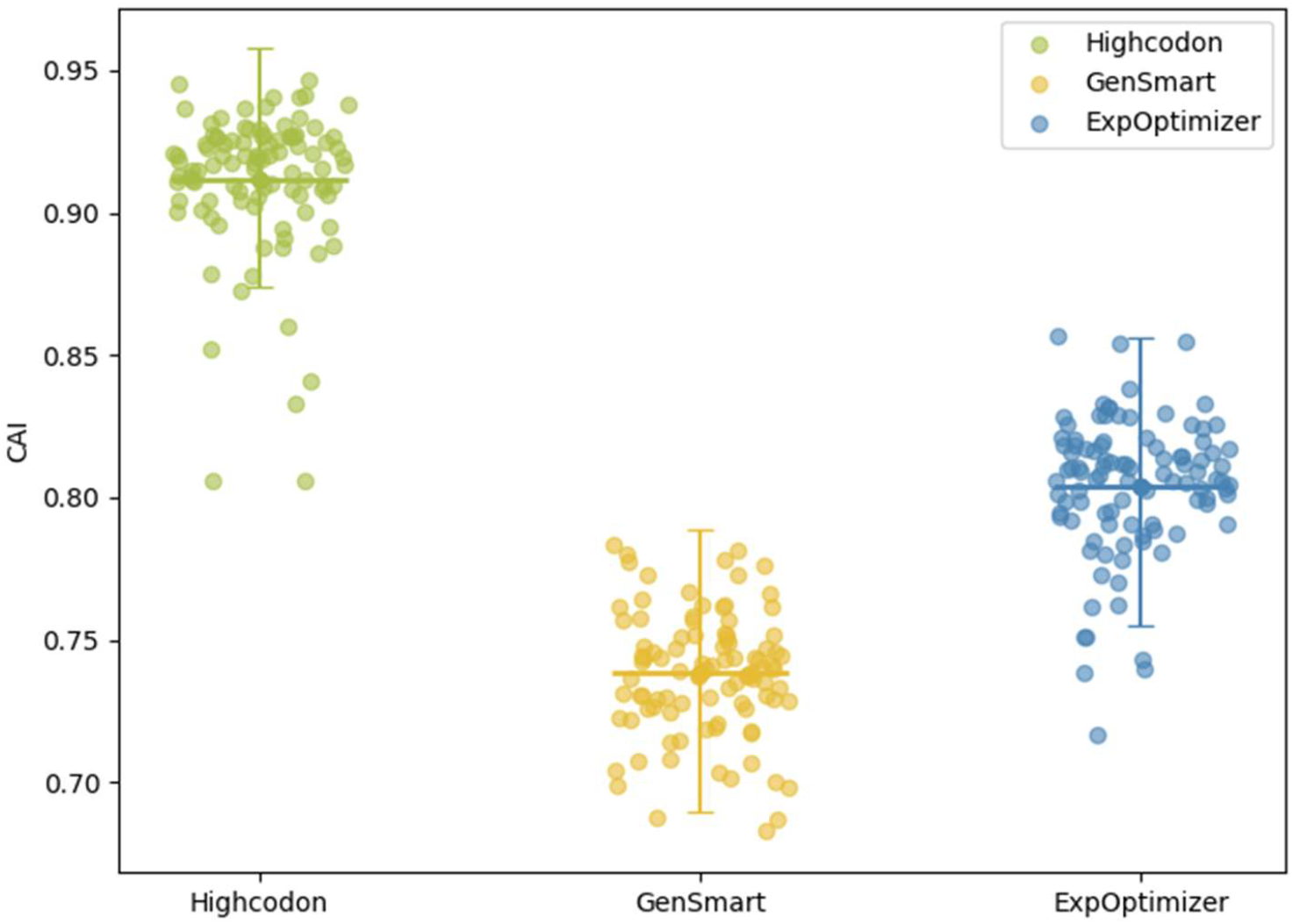

This figure shows the distribution of CAI values for three different tools: Highcodon, GenSmart, and ExpOptimizer. The x-axis represents the different models, and the y-axis indicates the CAI values. As can be seen from the figure, the CAI values of the Highcodon model are relatively concentrated and predominantly fall within the range of 0.85 to 0.95.Highcodon’s CAI value is approximately 23.45% higher than that of GenSmart and 13.34% higher than that of ExpOptimizer. Additionally, the error bar interval length of Highcodon is 15.12% shorter than that of GenSmart and 16.63% shorter than that of ExpOptimizer, further highlighting the stability of the Highcodon model,indicating the Highcodon model demonstrates higher stability and superiority in CAI values, which strongly supports its advantages in relevant research and applications.

### GC-content

**Figure 3.**
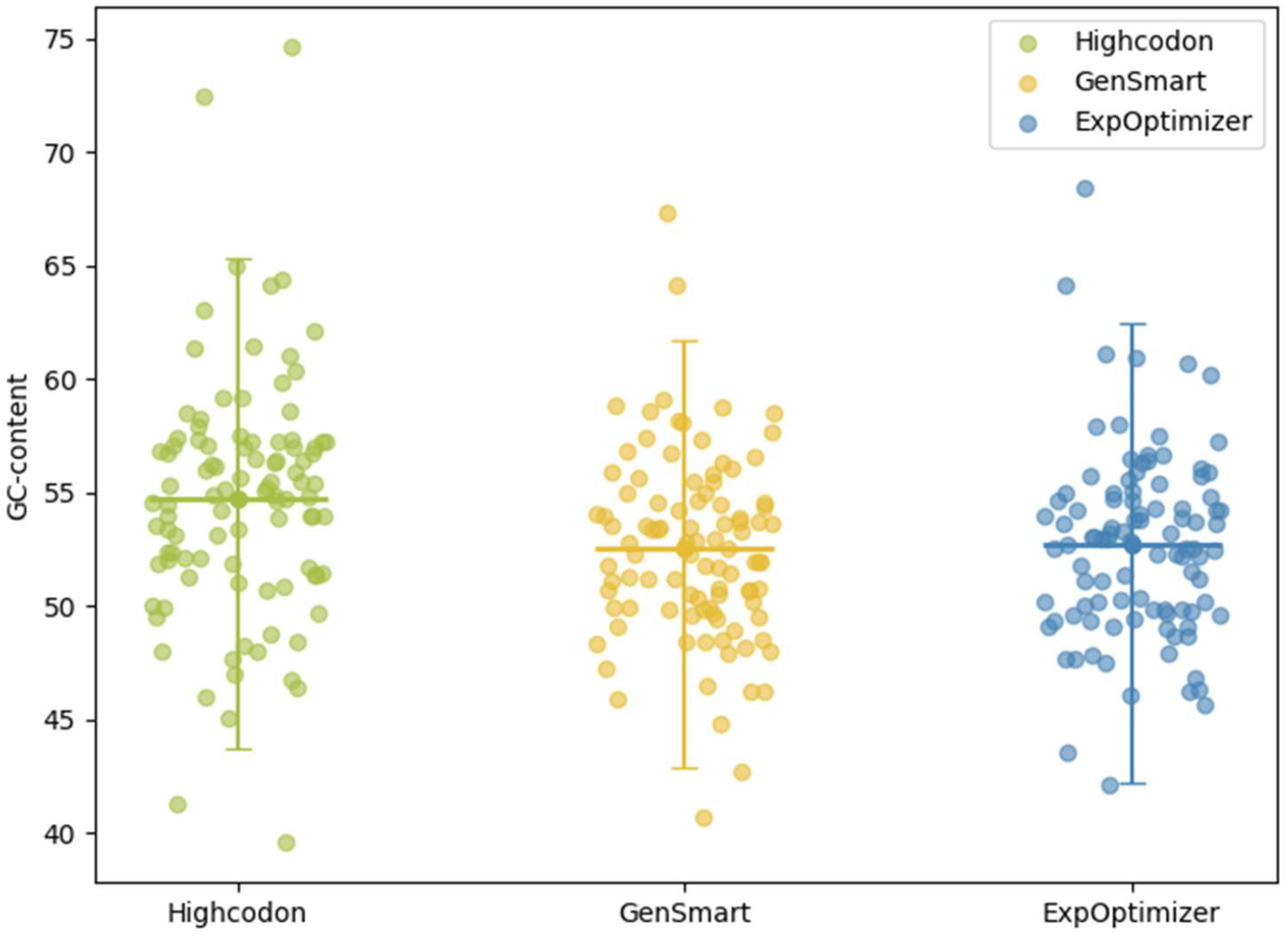

The GC content data points for Highcodon are primarily distributed between 45% and 60%, with a median value of approximately 55%. This indicates that in terms of GC content, Highcodon performs comparably to GenSmart and ExpOptimizer, as the majority of the GC content is concentrated within the optimal range.

## Methods

### Modeling

POS tagging is a fundamental task in natural language processing (NLP) that involves assigning the most appropriate part-of-speech tag to each word in a sentence^18^. Similarly, in the genetic coding system, there exists a many-to-one mapping between codons and amino acids. For example, leucine can be encoded by six different codons (TTA, TTG, CTT, CTC, CTA, CTG).

To address the codon optimization problem, we can formulate it as a sequence labeling task. In this framework, amino acid residues correspond to words in a sentence, while synonymous codons serve as the corresponding part-of-speech tags. Given an input amino acid sequence, the model assigns the optimal synonymous codon to each amino acid, ultimately generating a complete codon sequence. This analogy facilitates the application of NLP techniques to optimize codon selection in bioinformatics.

**Figure 4.**
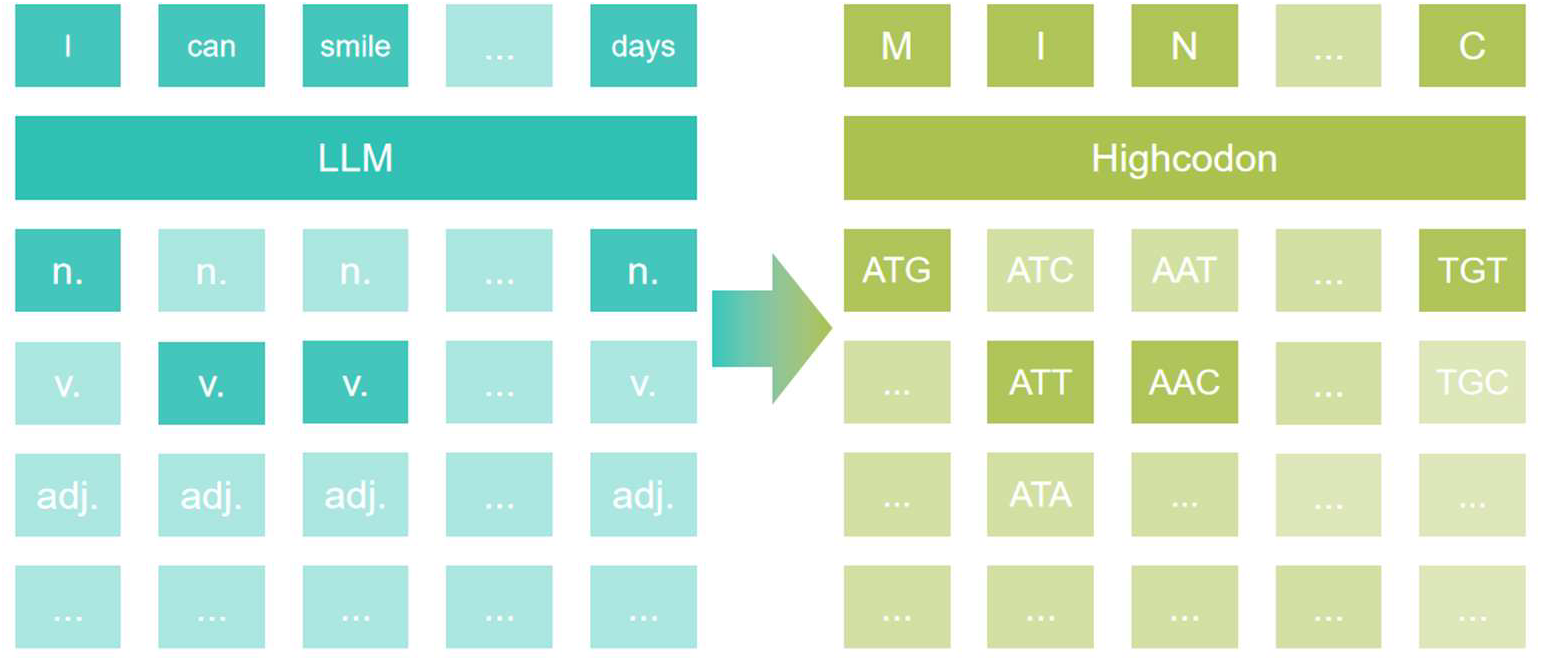

To achieve this goal, the model employs a pre-trained deep learning model based on BERT (Bidirectional Encoder Representations from Transformers), namely Prot_Bert, and fine-tunes it to perform a sequence labeling (token classification) task.

To better guide the model toward codon usage patterns associated with high protein expression, we introduced expression-weighted adjustments based on the standard cross-entropy loss function:

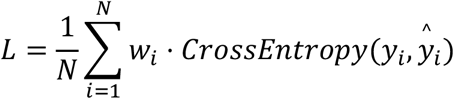

However, protein expression is not determined by codon usage alone, but rather results from a complex, non-linear interplay of multiple regulatory factors. Therefore, directly incorporating expression levels into the loss function may not be biologically rigorous. To address this, we adopted a more flexible strategy: we applied a tunable scaling factor *α*, and defined the final sample weight as:

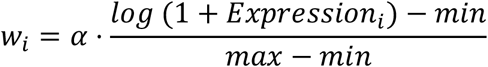

This formulation offers flexibility, as the influence of expression levels can be modulated by adjusting the scaling factor *α*, we can control the degree to which expression levels influence training.

This approach allows genes with higher expression levels to contribute more significantly during training, potentially steering the model toward codon preferences that are empirically linked to enhanced expression.

### Utilization of Pretrained Models

BERT (Bidirectional Encoder Representations from Transformers), proposed by Devlin et al. in 2018^19^, employs a bidirectional self-attention mechanism to enhance sequence understanding. In this study, we utilized the pretrained ProtBert model, which was unsupervisedly trained on the UniRef100 dataset containing 217 million protein sequences. During pretraining, 15% of amino acids were randomly masked: 80% were replaced with the [MASK] token, 10% with a random amino acid, and 10% remained unchanged. By leveraging the learned patterns of protein sequences, ProtBert could improve training efficiency and reduce computational costs compared to training from scratch

### Codon optimization

The goal of the Token classification task is that for an input sequence

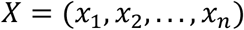

Model learning a mapping

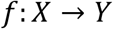

*Y* = (*y*_1_, *y*_2_, …, *y*_n_) is a codon tag sequence, *y*_i_ is the codon tag for amino acid *i* Therefore, in the BERT architecture, the output of the last Transformer layer *H* is taken as the token-level representation.

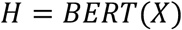

*H* = (*h*_1_, *h*_2_, …, *h*_n_) is the hidden state of each token

Add a fully connected layer on top of the BERT output to map the hidden states to the codon label space.

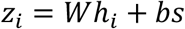

*W* and *b* are trainable parameters.

Finally, apply the Softmax activation function to compute the probability distribution of each token in the codon label space.

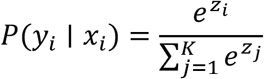

Under the constraint of biological plausibility, the model will select the codon with the highest probability as the final prediction.

**Figure 5.**
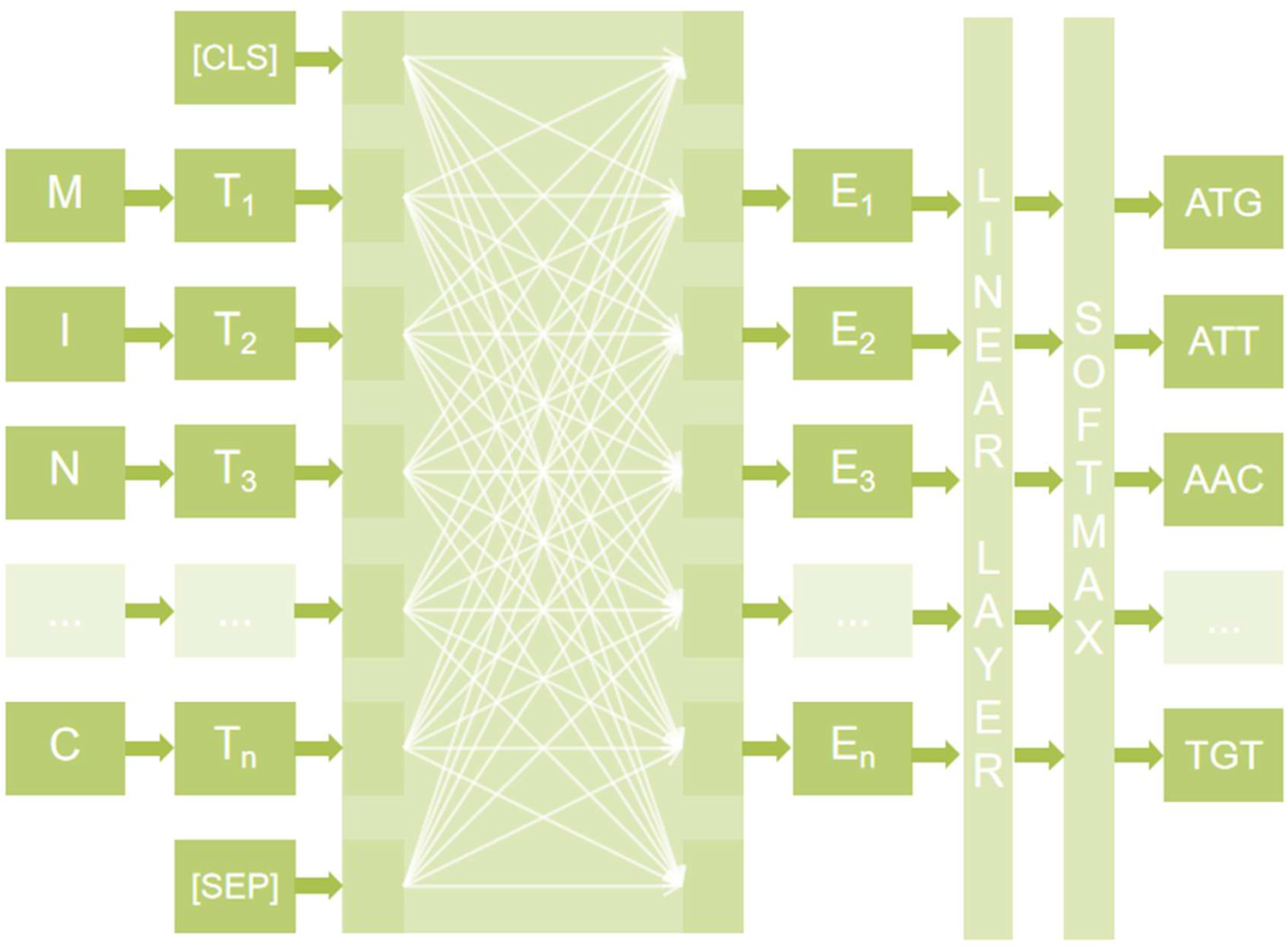

### Data

The expression data in this study were obtained from the PaxDb database (pax-db.org), specifically from the Escherichia coli - Whole Organism (Integrated) dataset. A total of 3,674 relative protein expression values for Escherichia coli were retrieved. The UniProtKB ID mapping tool was used to convert the dataset labels into GeneID format, and the corresponding gene sequences were downloaded from the National Center for Biotechnology Information (NCBI) database. After data processing, a total of 3,650 valid gene sequences were obtained.

The study conducted a statistical analysis of the length distribution of gene sequences. Given the architectural constraints of the BERT protein language model, which has a maximum processing sequence length of 512 amino acid residues, this study temporarily excluded DNA sequences exceeding 1,536 (512×3) nucleotides. This exclusion was implemented to optimize computational efficiency and reduce model complexity while ensuring biological information integrity. As a result, a total of 3,128 valid sequences were obtained.

## Protein expression

### Availability and requirements

The training data, validation data, training code, inference code, and the optimal model are available on GitHub (https://github.com/LIJIAWEI040301). The pre-trained model is Prot_BERT from Hugging Face (https://huggingface.co/Rostlab/prot_bert).

## Discussion

### Attention Analysis

The attention values increase along the diagonal, which aligns with fundamental principles, as codon usage is primarily influenced by the type of amino acid residues.

**Figure 6.**
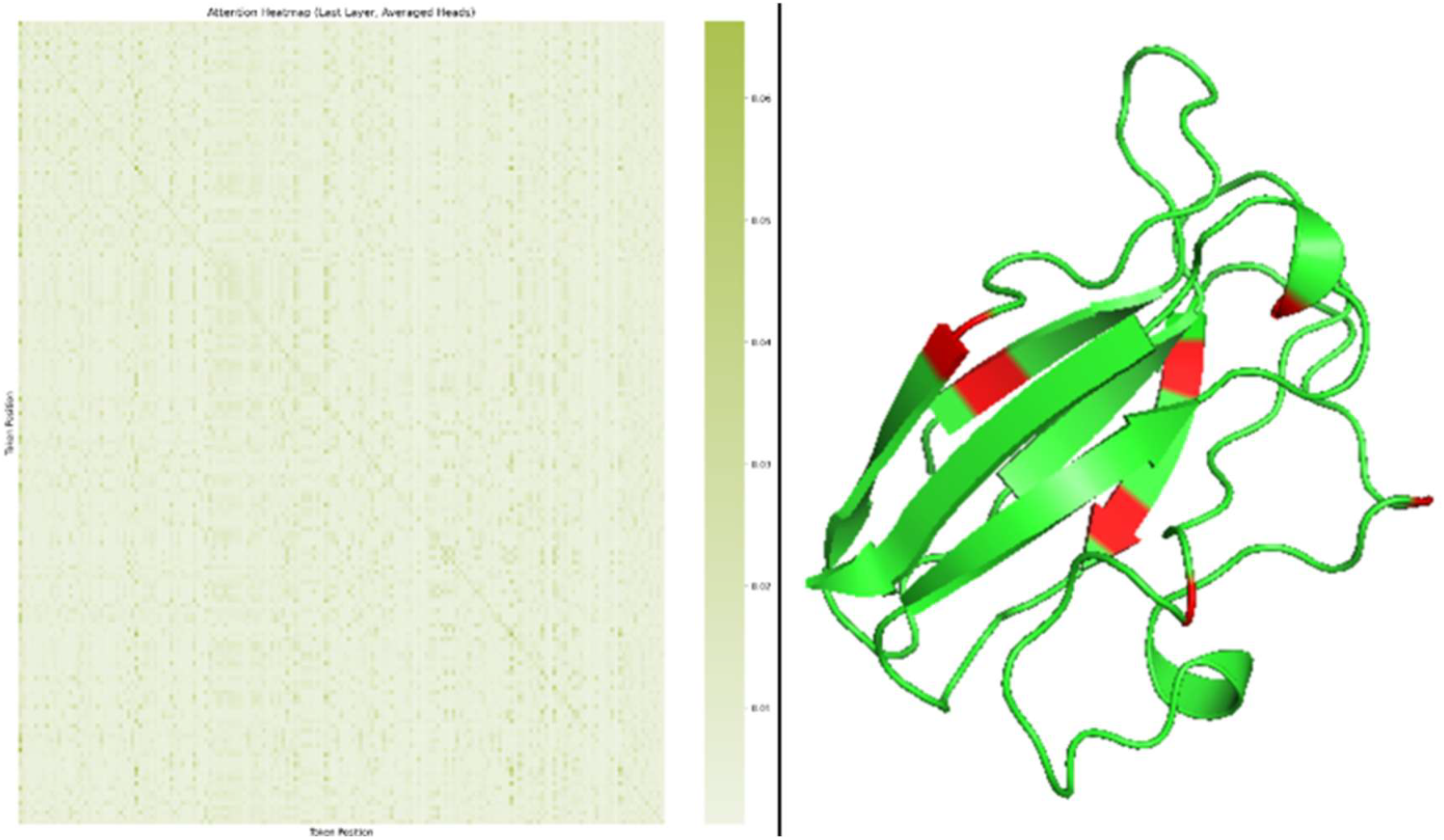

Furthermore, after modeling protein structures using AlphaFold^20^, we extracted high-confidence regions and mapped the top 10 positions with the highest attention scores. Interestingly, these high-attention regions were frequently located at structural turns, which are often associated with co-translational folding processes. Prior studies have suggested that excessive translation speed at such regions may interfere with proper folding, potentially resulting in structural abnormalities^21^.

These observations suggest that the model may have implicitly learned sequence features related to translation dynamics and folding-sensitive regions. While further experimental validation is required, this finding highlights the potential of attention-based models to uncover biologically relevant patterns beyond codon frequency alone. Such insights may inform future codon optimization strategies that go beyond simply selecting high-frequency or rapidly translated codons, particularly in structurally sensitive regions of the protein.

## Conclusion

In this study, we propose High-Codon, a deep learning-based method leveraging a BERT pre-trained model specifically designed for optimizing heterologous protein expression. High-Codon captures deeper codon usage patterns, including codon context dependency, and translation dynamics, enabling biologically informed optimization. Compared to existing methods, High-Codon exhibits greater applicability across different proteins and significantly enhances codon usage metrics associated with protein expression. We anticipate that High-Codon will provide a novel solution for efficient heterologous protein expression.

## Limitations and Challenges

Although the model has demonstrated strong performance, challenges remain in accurately quantifying high gene expression. While indirect metrics are avoided during feature selection, evaluation still relies on measures like the Codon Adaptation Index (CAI). Due to the triplet nature of codon usage, optimizing one metric may compromise another, making model selection experience-dependent. Prior knowledge of codon optimization can help balance these trade-offs. Thus, the model’s performance may not yet be fully optimized.

To improve, a linear regression model could predict gene expression from codon usage, reducing reliance on traditional metrics. Future work may include wet-lab experiments on selected test sequences to validate the optimization strategy and further integrate computational and experimental approaches.

## References

1. Khudainazarova, N. S. et al. Prokaryote- and Eukaryote-Based Expression Systems: Advances in Post-Pandemic Viral Antigen Production for Vaccines. IJMS 25, 11979 (2024).

2. Arbabi-Ghahroudi, M., Tanha, J. & MacKenzie, R. Prokaryotic expression of antibodies. Cancer Metastasis Rev 24, 501–519 (2005).

3. Yin, J., Li, G., Ren, X. & Herrler, G. Select what you need: A comparative evaluation of the advantages and limitations of frequently used expression systems for foreign genes. Journal of Biotechnology 127, 335–347 (2007).

4. Tuller, T., Waldman, Y. Y., Kupiec, M. & Ruppin, E. Translation efficiency is determined by both codon bias and folding energy. Proc. Natl. Acad. Sci. U.S.A. 107, 3645–3650 (2010).

5. Parvathy, S. T., Udayasuriyan, V. & Bhadana, V. Codon usage bias. Mol Biol Rep 49, 539–565 (2022).

6. Hershberg, R. & Petrov, D. A. General Rules for Optimal Codon Choice. PLoS Genet 5, e1000556 (2009).

7. Gouy, M. & Gautier, C. Codon usage in bacteria: correlation with gene expressivity. Nucl Acids Res 10, 7055–7074 (1982).

8. Hershberg, R. & Petrov, D. A. Selection on Codon Bias. Annu. Rev. Genet. 42, 287–299 (2008).

9. Gustafsson, C., Govindarajan, S. & Minshull, J. Codon bias and heterologous protein expression. Trends in Biotechnology 22, 346–353 (2004).

10. Villalobos, A., Ness, J. E., Gustafsson, C., Minshull, J. & Govindarajan, S. Gene Designer: a synthetic biology tool for constructing artificial DNA segments. BMC Bioinformatics 7, 285 (2006).

11. Kudla, G., Murray, A. W., Tollervey, D. & Plotkin, J. B. Coding-Sequence Determinants of Gene Expression in Escherichia coli. Science 324, 255–258 (2009).

12. Liu, Y. A code within the genetic code: codon usage regulates co-translational protein folding. Cell Commun Signal 18, 145 (2020).

13. Fu, H. et al. Codon optimization with deep learning to enhance protein expression. Sci Rep 10, 17617 (2020).

14. Jain, R., Jain, A., Mauro, E., LeShane, K. & Densmore, D. ICOR: improving codon optimization with recurrent neural networks. BMC Bioinformatics 24, 132 (2023).

15. Gong, H. et al. Integrated mRNA sequence optimization using deep learning. Briefings in Bioinformatics 24, bbad001 (2023).

16. Goulet, D. R. et al. Codon Optimization Using a Recurrent Neural Network. Journal of Computational Biology 30, 70–81 (2023).

17. Sharp, P. M. & Li, W.-H. The codon adaptation index-a measure of directional synonymous codon usage bias, and its potential applications. Nucl Acids Res 15, 1281–1295 (1987).

18. Chiche, A. & Yitagesu, B. Part of speech tagging: a systematic review of deep learning and machine learning approaches. J Big Data 9, 10 (2022).

19. Devlin, J., Chang, M.-W., Lee, K. & Toutanova, K. BERT: Pre-training of Deep Bidirectional Transformers for Language Understanding. Preprint at http://arxiv.org/abs/1810.04805 (2019).

20. Jumper, J. et al. Highly accurate protein structure prediction with AlphaFold. Nature 596, 583–589 (2021).

21. Perach, M., Zafrir, Z., Tuller, T. & Lewinson, O. Identification of conserved slow codons that are important for protein expression and function. RNA Biology 18, 2296– 2307 (2021).

